# Stochastic Modeling and Transient Analysis of Epidemic Extinction Dynamics

**DOI:** 10.1101/2025.10.13.682061

**Authors:** Baoyin Yuan, Feng Jiao

**Affiliations:** School of Mathematics and Information Science, Guangzhou University, Guangzhou 510006, People’s Republic of China; Guangzhou Center for Applied Mathematics, Guangzhou University, Guangzhou 510006, People’s Republic of China

**Keywords:** Generation time, finite-time extinction, branching process, renewal equation, probability generating function

## Abstract

Understanding extinction probabilities in branching processes is pivotal for epidemiology and population dynamics. Traditional models often assume a fixed generation time, resulting in extinction probabilities determined solely by offspring distributions and remaining unchanged over time. By contrast, our study incorporates the generation time into the analysis, considering how the timing of each generation influences extinction dynamics. By focusing on the finite-time risk of extinction, our approach reveals that shorter generation times can lead to a temporary increase in the risk of population or epidemic die-out. We support our findings with precise fixed-point analyses and numerical integration techniques based on real data from various infectious diseases. Although the long-term probability of extinction does not change, the transient dynamics show that a faster-paced transmission process may elevate early extinction risk. The study highlights the crucial role of transmission timing in epidemic modeling and indicates that accounting for generation time can provide new perspectives for developing effective public health strategies and outbreak control measures.

## 1 Introduction

In epidemiology and population dynamics, understanding the likelihood that a population eventually dies out is essential for predicting outbreaks and managing species survival. Mathematical modeling provides a systematic way to represent complex biological processes in a simplified form (Anderson and May 1991). By using equations and computational simulations, models enable researchers to capture key dynamics, investigate the effects of different parameters, and make predictions about future behavior. This approach has proven invaluable for elucidating the mechanisms behind epidemic spread and population changes, ultimately guiding public health decisions and conservation efforts (Kucharski et al. 2020; Sun et al. 2021; Wu et al. 2020).

At its core, a branching process is a mathematical framework that describes population evolution, where each individual produces a random number of offspring according to a specified probability distribution. In essence, the process decomposes population dynamics into fundamental “branching” steps. In branching processes, the probability that a population eventually dies out (known as the extinction probability) is a central topic in mathematical biology, with applications ranging from population dynamics to the spread of infectious diseases. Harris (Harris 1963) and Athreya and Ney (Athreya and Ney 1972) laid the groundwork by identifying the offspring distribution as the primary determinant of extinction probability. Offspring distribution quantifies the likelihood of an individual producing a given number of offspring and is mathematically expressed through the probability generating function (PGF)

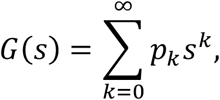

where *p*_*k*_ denotes the probability that an individual produces exactly *k* offspring and *s* is a placeholder variable that allows the offspring distribution to be encoded compactly. The eventual extinction probability, denoted by *q*(∞), (i.e., *q* in short), is then obtained as the smallest solution in [0,1] to the fixed-point equation

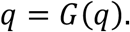

These classical models assume that each individual produces offspring following a predefined distribution. Common choices include the Poisson distribution (appropriate when events occur randomly and independently) and the geometric distribution (suitable when the probability of an event remains constant over time). A key parameter is the basic reproduction number, *R*_0_, which represents the average number of offspring (or secondary infections) produced by one individual. When *R*_0_ ≤ 1, extinction is almost certain, while if *R*_0_ > 1, there exists a nonzero chance of indefinite persistence (Harris 1963; Athreya and Ney 1972; Mode 1971; Becker 1989).

Subsequent extensions of these ideas have incorporated heterogeneity in the offspring distribution. Lloyd-Smith et al. (2005) highlighted the importance of individual variation and superspreading events in infectious disease dynamics, demonstrating that variability in the number of secondary cases can significantly alter the emergence and persistence of epidemics. Keeling and Rohani (2008) and Anderson and May (1991) emphasized that higher moments of the offspring distribution (e.g., variance and skewness) can have a significant impact on extinction risks (Brauer and CastilloChavez 2012;Wallinga and Lipsitch 2007).

Despite the extensive body of research on extinction probabilities and offspring distributions (Becker 1989; Lloyd 2001; Nishiura 2009; Lipsitch et al. 2003; Ferguson 2006), most classical models treat extinction probability as a stationary, time-independent quantity. Consequently, these studies focus primarily on the long-term probability of extinction. However, in many practical scenarios—particularly during the early stages of an outbreak or amid rapid ecological disturbances—a time-dependent understanding of extinction dynamics is essential. Decision-makers may require insights into the transient behavior of extinction probabilities, i.e., the probability of extinction within a finite time horizon rather than solely its asymptotic limit.

One critical yet underexplored aspect in this context is the role of the generation time distribution, which characterizes the interval between successive reproduction or transmission events. Early models often assumed a constant generation time or neglected its variability entirely. Empirical studies ( Svensson 2007; Wallinga and Lipsitch 2007) have since demonstrated that the generation time (which may include incubation periods and latency effects) significantly affects outbreak dynamics. For instance, realistic generation time distributions modeled using exponential or gamma functions have been shown to shape transient epidemic trajectories in a manner that can critically influence intervention strategies (Fisher 1992; Feehan and Mahmud 2021; He et al. 2020).

Recent investigations into the COVID-19 pandemic (He et al. 2020; Ferretti et al. 2020; Chinazzi et al. 2020; Kucharski et al. 2020; Park et al. 2020) have underscored the critical impact of generation time variability on transmission potential and epidemic curves. These studies indicate that neglecting the generation time distribution can lead to underestimation or oversimplification of early epidemic behavior (Adam et al. 2020; Ma 2020). In addition, the interplay between transmission heterogeneities and generation time parameters has been investigated across several in multiple recent analytical and computational frameworks (Wu et al. 2020; Brauner and Castillo-Chavez 2021; Sun et al. 2021). Such approaches offer a refined understanding of outbreak dynamics, particularly valuable for implementing adaptive intervention policies (Ward et al. 2023; Diekmann et al. 1990).

Motivated by these developments, the present study bridges the gap between classical time-invariant models and the need to understand finite-time dynamics by incorporating the generation time distribution into the branching process framework. To measure the risk that a population will die out within a specified period, we introduce the concept of a finite-time extinction probability *q*(*t*; *T*_*g*_), with *T*_*g*_ representing the mean generation time and *t* being the calendar time. Our mathematical analysis demonstrates that *q*(*t*; *T*_*g*_) is negatively correlated with *T*_*g*_. In other words, shorter generation times may transiently increase the extinction risk, even if the asymptotic extinction probability remains unchanged. We validate these theoretical findings through numerical simulations that generate an array of visualizations (including time-series plots, fixed-time scans, and multidimensional contours) to elucidate the interplay between extinction probability and generation time.

The remainder of this study is organized as follows. Section 2 derives the finite-time extinction probability equation in the context of infectious disease transmission, establishing the foundational mathematical framework for our analysis. Section 3 presents a detailed investigation into the relationship between the finite-time extinction probability, *q*(*t*; *T*_*g*_), and the mean generation time, *T*_*g*_, demonstrating a precise and quantifiable connection between these two key parameters. Section 4 employs numerical simulations to examine the evolution of *q*(*t*; *T*_*g*_) as a function of the mean generation time and the combined effects of the generation time and offspring distributions on the extinction process. Section 5 discusses the implications of our findings for public health and epidemic modeling, and outlines potential future extensions. Finally, Section 6 provides detailed technical information on the numerical simulation methods used throughout the study.

## 2 Modeling Finite-Time Extinction Probability in Disease Transmission

We derive the finite-time extinction probability equation in the context of infectious disease transmission. Subsequently, the relationship between finite-time extinction probability and the mean generation time is analyzed in a combination of mathematical derivation and numerical simulations in the subsequent sections.

### 2.1 Biological and Mathematical Context

In an infectious disease outbreak, an infected individual generates a number of secondary infections following some offspring distribution. Let *p*_*k*_ denote the probability that an individual produces exactly *k* secondary cases. The classical PGF is given by 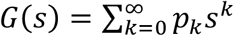, and the ultimate extinction probability is obtained as the smallest solution of *q* = *G*(*q*).

In real epidemics, however, public health decisions often require an understanding of the extinction probability within a finite time frame. Biological factors such as incubation periods, delays in symptom onset, and inherent reproduction delays (the generation time) are critical. To capture these effects, we assume that the delay (or generation time) between an individual’s infection and the production of secondary infections is a random variable with probability density function *f*_*τ*_(*t*) and cumulative distribution function (CDF)

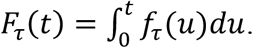

For instance, the generation time can be modeled using an exponential distribution, expressed by the following equation:

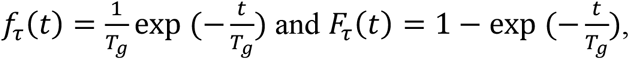

where *T*_*g*_ denotes the mean generation time. Biologically, *F*_*τ*_(*t*) represents the probability that an individual has produced secondary cases by time *t*.

### 2.2 Derivation of the Finite-Time Extinction Probability Equation

To derive the finite-time extinction probability, consider the two mutually exclusive scenarios for an infected individual within a finite time *t*. To represent the CDF that explicitly incorporates the dynamics imposed by the generation time *T*_*g*_ into account, we extend the notation *F*_*τ*_(*t*) to *F*_*τ*_(*t*; *T*_*g*_), which describes the probability that, given a generation time *T*_*g*_, an individual has produced secondary cases by time *t*.

**Case 1:** The infected individual does not produce any secondary infections within time *t*. The probability of this event is given by the product of two factor: the probability of not reproducing when the opportunity is present, *p*_0_, and the probability that no reproduction occurs due to the generation time delay 1 − *F*_*τ*_(*t*; *T*_*g*_). This component is expressed as follows:

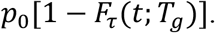

**Case 2:** The individual produces secondary cases at some time *τ* ∈ [0, *t*], after which those secondary cases must go extinct within the remaining time *t* − *τ*. The probability that reproduction occurs at time *τ* is given by *f*_*τ*_(*τ*; *T*_*g*_), while the extinction of the resulting lineage is described by the PGF evaluated at the extinction probability over the remaining time-window, i.e., *G*(*q*(*t* − *τ*; *T*_*g*_)). Integrating over all possible *τ* from 0 to *t*, the contribution is given as follows:

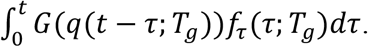

Thus, by combining the two components, the finite-time extinction probability is given by the following integral equation:

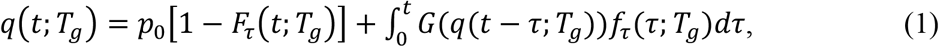

with the initial condition *q*(0; *T*_*g*_) = *p*_0_, since there is no time for reproduction at *t* = 0.

By incorporating a time delay, this equation extends the classical fixed-point formulation and integrates key biological processes, such as incubation periods and transmission delays, directly into the extinction probability model.

## 3 Analytical Examination: Relationship with Mean Generation Time

The average generation time, *T*_*g*_, reflects the delay between an infection and when an individual begins transmitting the disease. Shorter generation times imply faster transmission, which, within a finite time window, can lead to an increased turnover of infections. Paradoxically, rapid turnover may increase the finite-time extinction probability because many potential transmission events are pruned either by early interventions or natural recovery before they effectively contribute to sustained transmission.

Our analysis establishes a precise relationship between the finite-time extinction probability *q*(*t*; *T*_*g*_)and the mean generation time *T*_*g*_. In particular, we show that the sensitivity of *q*(*t*; *T*_*g*_) with respect to *T*_*g*_ is governed by a Volterra-type integral equation, and that—under standard assumptions—this sensitivity is strictly negative.

### 3.1 Scale-Family Transformation

Under the scale-family assumption for the generation time distribution, there exist a standard CDF *F*_0_(*u*) and a corresponding density *f*_0_(*u*), defined as follows:

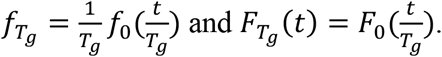

By substituting these representations into the original renewal equation for the finite-time extinction probability, and applying the change of variables *u* = *τ*/*T*_*g*_ (with *τ* = *T*_*g*_*u* and *dτ* = *T*_*g*_*du*), we obtain an equivalent formulation as follows:

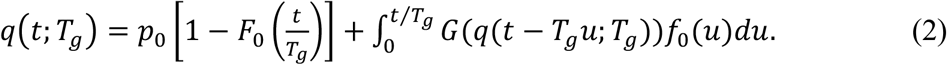

This representation effectively decouples the time variable from the scale parameter *T*_*g*_, providing a streamlined foundation for the subsequent sensitivity analysis.

### 3.2 Derivation of a Volterra-Type Integral Equation for the *T*_*g*_ Derivative

We begin by defining the sensitivity function as follows:

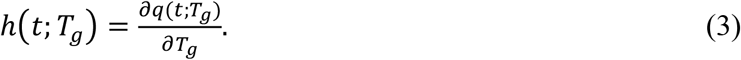

This function represents the derivative of the finite-time extinction probability with respect to the mean generation time *T*_*g*_. In **Appendix A**, we rigorously prove the differentiability of *q*(*t*; *T*_*g*_) with respect to *T*_*g*_. To ensure that the derivation is valid and mathematically comprehensive, the following assumptions are required:

**Assumptions on Smoothness and Differentiability:**

1. The standard density *f*_0_(*u*), which underlies the generation time distribution through the scale-family transformation, is continuously differentiable, bounded, and strictly positive on its support.
2. The generating function *G*(*s*), which characterizes the offspring distribution, is strictly increasing, continuously differentiable, and bounded on the interval [0,1], with its derivative *G*^′^(*s*) bounded.

The first assumption ensures that operations such as the change in variables (e.g., *u* = *τ*/*T*_*g*_) and the subsequent integration are well defined and avoid pathological behaviors; the second assumption guarantees the existence of all necessary derivatives and justifies the use of the chain rule and Leibniz’s integral rule when differentiating the renewal equation. Overall, these assumptions provide a solid analytical foundation for defining and deriving the associated Volterra-type integral equation as follows.

#### Lemma 3.1

Let *q*(*t*; *T*_*g*_) be expressed in the scale-transformed form described above. Subsequently, the sensitivity function

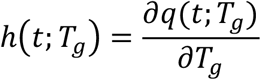

satisfies the following integral equation:

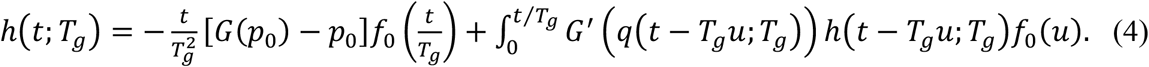

Intuitively, the first term on the right-hand side arises from differentiating the “delayed reproduction” (i.e., nonreproduction) component—capturing the effect that if an individual does not have the opportunity to reproduce before time *t*, this contribution is sensitive to *T*_*g*_. The second term, obtained via Leibniz’s rule, aggregates the contributions from all reproduction events that occur prior to time *t*. For a complete derivation, including the application of the chain rule and the Leibniz integral rule, please refer to **Appendix B**.

### 3.3 Monotonicity of *q*(*t*; *T*_*g*_) with Respect to *T*_*g*_

Building on **Lemma 3.1**, we now address the monotonicity of the finite-time extinction probability with respect to *T*_*g*_. Our primary result is stated as follows.

#### Theorem 3.2

Let the generating function

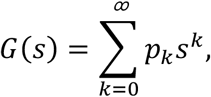

satisfy the following conditions:

1. Assumptions on Smoothness and Differentiability about *G*(*s*) and *f*_0_(*u*).
2. Nontrivial Offspring Distribution: The offspring distribution is nontrivial, meaning that *p*_0_ < 1 and there exists at least one *k* ≥ 1 such that *p*_*k*_ > 0. This condition ensures, in particular, that 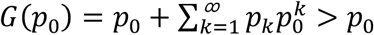

Subsequently, for every *t* > 0, the finite-time extinction probability *q*(*t*; *T*_*g*_) is strictly decreasing as a function of the mean generation time *T*_*g*_; i.e.,

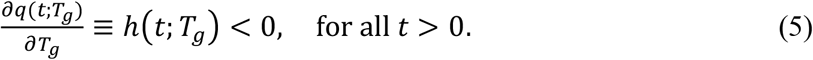

The proof proceeds by introducing an auxiliary function *φ*(*t*) = −*h*(*t*; *T*_*g*_), which satisfies a similar Volterra-type integral equation:

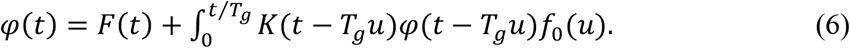

with

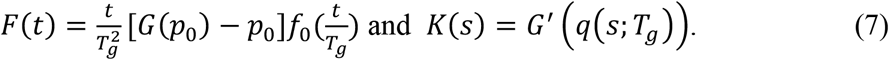

Since *F*(*t*) > 0 and *K*(*s*) > 0, a fixed-point iterative procedure (detailed in **Appendix C**) shows that *φ*(*t*) > 0 for all *t* > 0. Consequently, we deduce that *h*(*t*; *T*_*g*_) = −*φ*(*t*) < 0, thereby establishing the desired monotonicity.

In the remainder of the study, we provide a detailed discussion of the assumptions required for our analysis and outline the implications of our findings in the context of infectious disease transmission. The complete derivations and proofs of **Lemma** and **Theorem** are included in **Appendixes B** and **C**, respectively, where all key technical steps are presented.

## 4 Numerical Simulation

In this section, we employ numerical simulations to investigate: (1) how the finite-time extinction probability *q*(*t*; *T*_*g*_) evolves with time and with the mean generation time *T*_*g*_ and (2) how the interplay between generation time and offspring distribution influences the extinction progress. The model applies to infectious disease transmission scenarios and broader biological population dynamics, where the stochasticity in individual reproduction plays a central role.

### 4.1 Evolution of the Finite-Time Extinction Probability

Our numerical simulation produced several insightful outcomes regarding the finite-time extinction probability *q*(*t*; *T*_*g*_) as a function of time *t* and the mean generation time *T*_*g*_. As shown in **Fig. 1**, for a fixed basic reproduction number *R*_0_ = 1.5, we observe that the finite-time extinction probability increases with time and eventually converges to an asymptotic value *q*(∞). In scenarios with a shorter mean generation time, the extinction probability rises more rapidly, indicating that although a faster transmission of infection may lead to an early surge in cases, it concurrently tends to drive the process toward extinction at an earlier stage.

**Figure 1.**
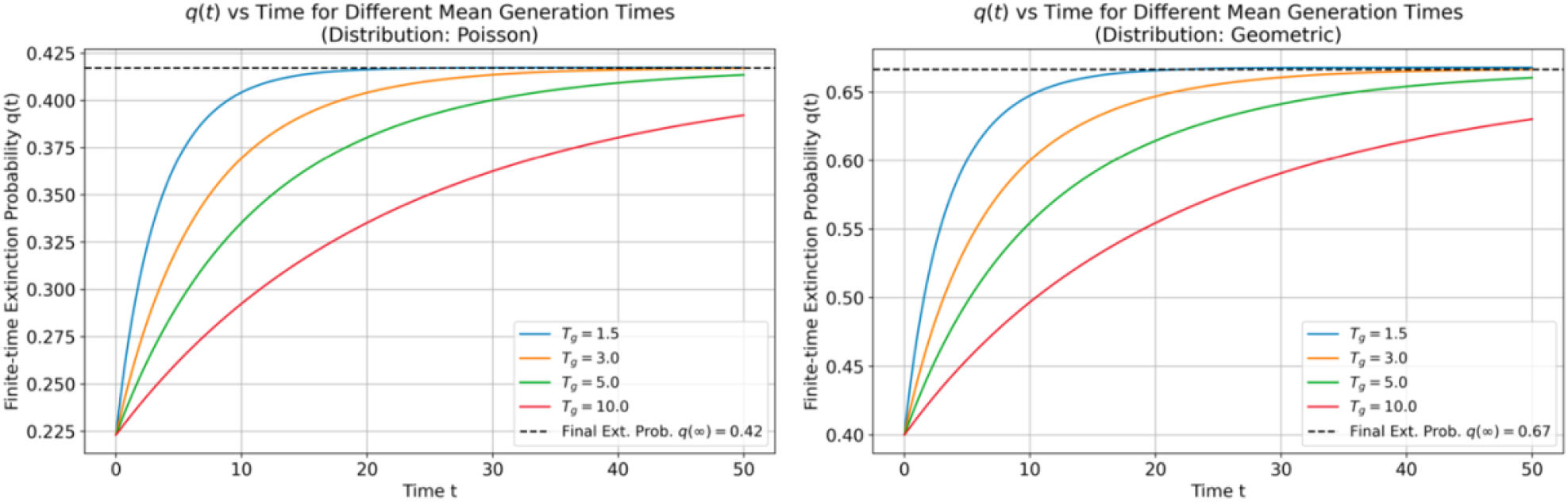
Time evolution curves of the finite-time extinction probability *q*(*t*; *T*_*g*_) as a function of time *t* for various mean generation times *T*_*g*_, with a fixed basic reproduction number *R*_0_ = 1.5. The figure illustrates how shorter generation times lead to a more rapid convergence toward the asymptotic extinction probability *q*(∞).

In addition, when examining the model through parameter scans as in **Fig. 2**, where *q*(*t*; *T*_*g*_) is evaluated at fixed time points (such as *t* = 2,5, and 8), it becomes evident that the probability of extinction decreases smoothly as *T*_*g*_ increases. This suggests that longer generation times, which effectively delay transmission events, reduce the likelihood of rapid extinction during the early phases of the process. To further elucidate these dynamics by highlighting the interplay between time *t* and *T*_*g*_, the heatmaps in **Fig. 3** are plotted to exhibit a clear color gradient corresponding to changes in extinction probability.

**Figure 2.**
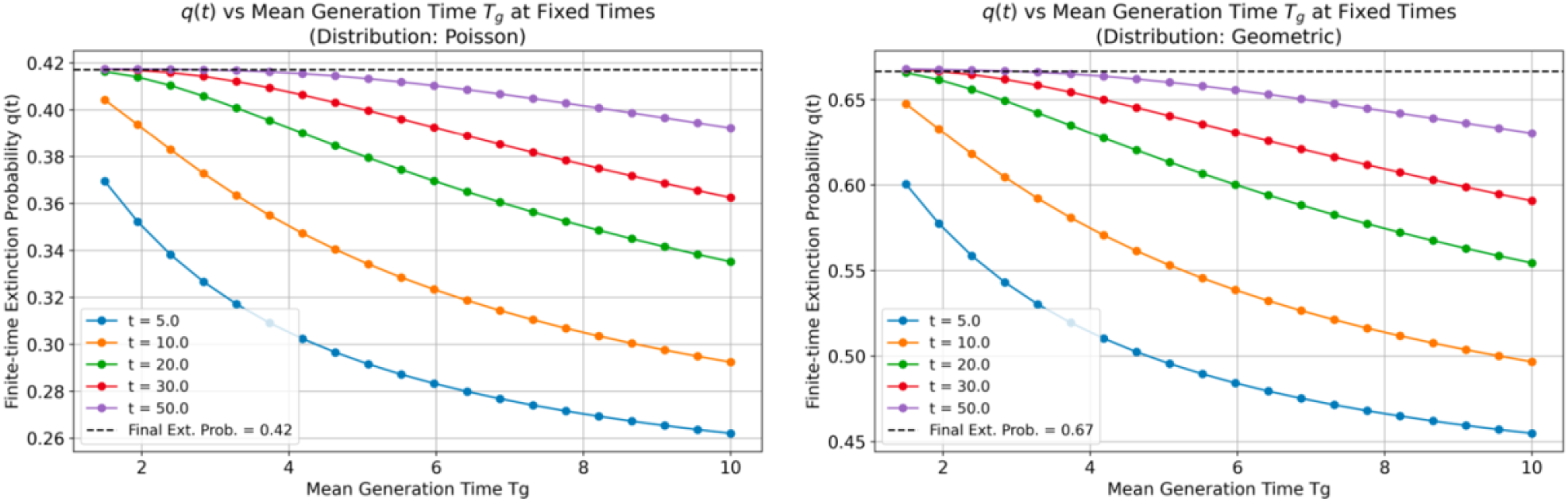
Parameter scan plots showing *q*(*t*; *T*_*g*_) at fixed time points (e.g., *t* = 5, 10, 20, 30, and 50) across a range of *T*_*g*_ values, illustrating the sensitivity of extinction probability to changes in mean generation time.

**Figure 3.**
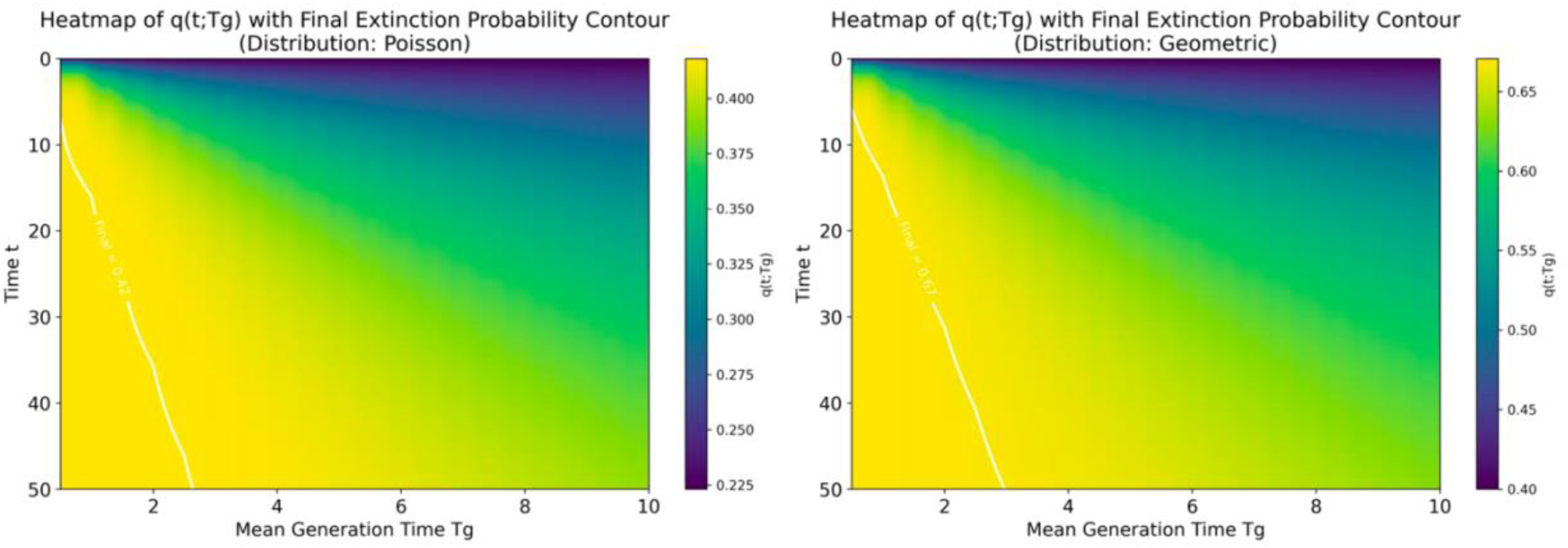
Heatmap of *q*(*t*; *T*_*g*_) across the time range and a spectrum of mean generation times, with a color gradient indicating extinction probabilities and overlaid contour lines representing the asymptotic value *q*(∞).

Poisson and geometric distributions are used to characterize the offspring distribution using only the single parameter *R*_0_, thereby ignoring any variation in offspring numbers. When comparing **Figs. 1-3** that depict the finite-time extinction probability *q*(*t*; *T*_*g*_) under a reproduction number of 1.5, both distributions yield generally similar evolutionary patterns. The only notable difference is that the geometric distribution results in a higher final extinction probability and, correspondingly, higher values of *q*(*t*; *T*_*g*_) at finite times.

### 4.2 Impact of Offspring Variability on Finite-Time Extinction Risk

In epidemic modeling, the negative binomial distribution is often utilized to capture individual-level heterogeneity in the number of secondary cases produced by an infected individual (Lloyd-Smith et al. 2005). The dispersion parameter *k* within this distribution quantifies the degree of variation in transmission. A smaller *k* indicates higher variability, meaning that while most infected individuals might generate only a few secondary cases, a few “superspreaders” can be responsible for a disproportionately large number of infections. Conversely, as *k* ⟶ ∞, the negative binomial distribution converges to the Poisson distribution, where the variation in the number of secondary cases is minimal, and when *k* = 1, it becomes equivalent to the geometric, reflecting a moderate level of dispersion.

**Fig. 4** shows the temporal evolution of the finite time extinction probability *q*(*t*; *T*_*g*_) for each of the nine infectious diseases: SARS, measles, MERS, smallpox, pneumonic plague, COVID-19, A/H1N1, hantavirus, and Ebola, the parameters of which are listed in **Table 1**. A key observation from this figure is that, despite following different trajectories, all diseases eventually converge to their respective asymptotic extinction probabilities *q*(∞). However, the paths to convergence vary markedly. For instance, measles—characterized by the highest *R*_0_ = 15 — reaches a near-steady state rapidly, indicating that an inevitable outbreak occurs almost immediately. By contrast, for SARS and MERS, where *R*_0_ < 1, the evolution of *q*(*t*; *T*_*g*_) is markedly different. These differences can be attributed to variations in the dispersion parameter *k* and the mean generation time *T*_*g*_. The trajectories of different diseases may intersect at various time points despite their definite distinct limiting *q*(∞) values. This observation underscores the importance of the definition of finite-time extinction probability when analyzing epidemic dynamics. For instance, while the eventual extinction probability of MERS is higher than that of pneumonic plague, there exist intermediate time points at which the instantaneous extinction probability for Pneumonic Plague may be lower than that for MERS. This distinction is crucial for understanding the nuances of epidemic evolution and can have significant implications for public health interventions.

**Table 1.** Basic data of different infectious diseases and the references for the data source.

| Infectious disease | Epidemiological parameters<br>($R_0$: the basic reproduction number; $k$:<br>the dispersion parameter; $T_g$: the mean<br>generation time) | References |
| --- | --- | --- |
| SARS | $R_0 = 1.63, k = 0.16, T_g = 8.4\text{days}$ | Lloyd-Smith et al. 2005;<br>Wallinga and Teunis 2004 |
| Measles | $R_0 = 9.2, k = 0.32, T_g = 10.7\text{days}$ | Nishiura et al. 2017; Vink et<br>al. 2014 |
| MERS | $R_0 = 0.91, k = 0.06, T_g = 10.7\text{days}$ | Cauchemez et al 2014;<br>Chowell et al. 2015 |
| Smallpox | $R_0 = 3.19, k = 0.37, T_g = 16.0\text{days}$ | Lloyd-Smith et al. 2005;<br>Nishiura and Eichner 2007 |
| Pneumonic Plague | $R_0 = 1.32, k = 1.37, T_g = 4.86\text{days}$ | Lloyd-Smith et al. 2005;<br>Nishiura et al. 2006 |
| COVID-19 | $R_0 = 2.5, k = 0.1, T_g = 4.21\text{days}$ | Chen 2022; Endo et al. 2020 |
| A/H1N1 | $R_0 = 1.2, k = 0.94, T_g = 3.0\text{days}$ | Boëlle et al. 2011; Fraser et<br>al. 2009 |
| Hantavirus | $R_0 = 0.70, k = 1.66, T_g = 18.0\text{days}$ | Lloyd-Smith et al. 2005;<br>Vial et al. 2006 |
| Ebola | $R_0 = 1.5, k = 5.1, T_g = 14.0\text{days}$ | Lloyd-Smith et al. 2005;<br>Van Kerkhove et al. 2015 |

**Figure 4.**
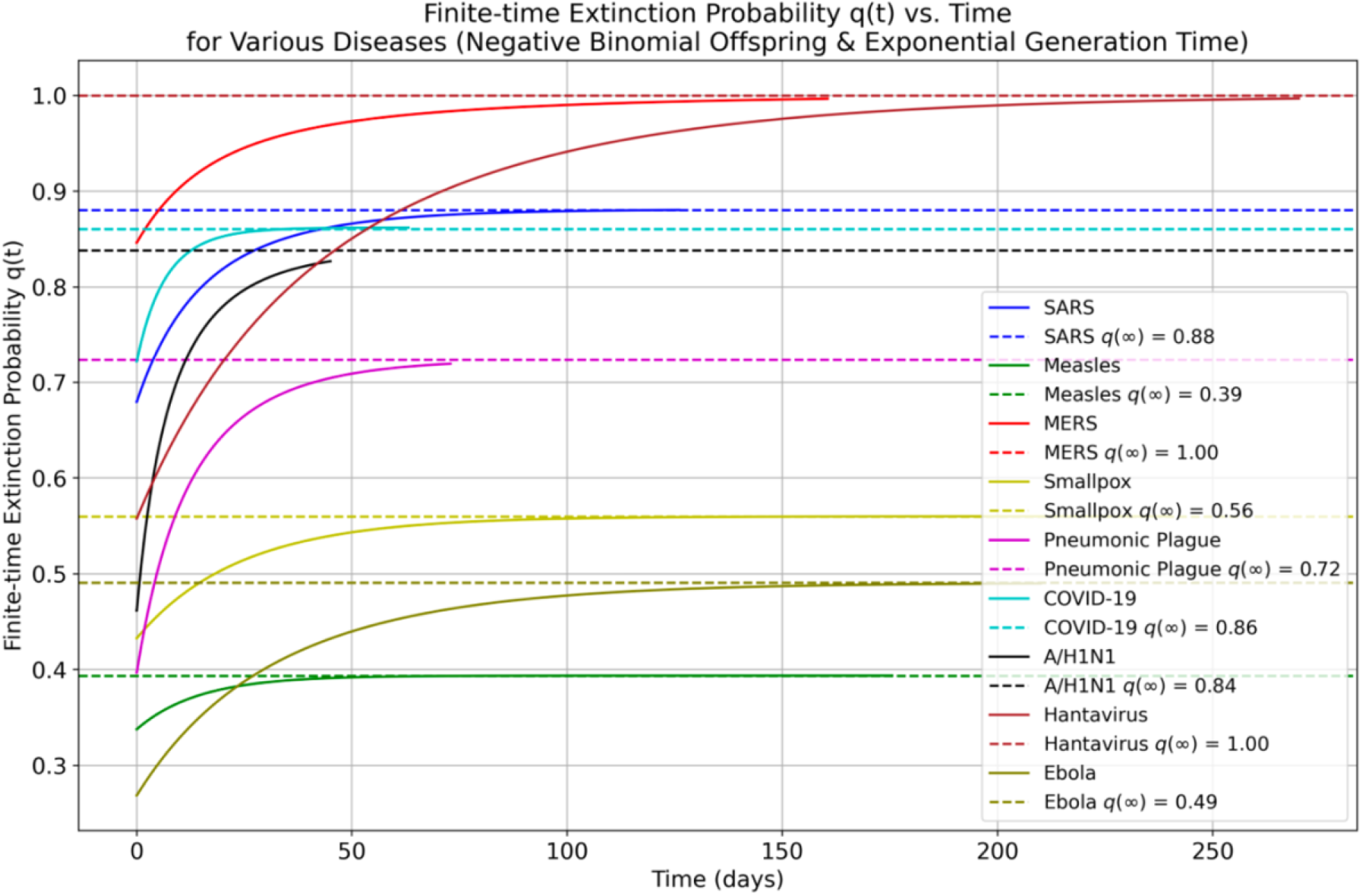
Trajectory of the finite-time extinction probability *q*(*t*; *T*_*g*_) approaching its respective asymptotic value *q*(∞) over time. Each of the nine infectious diseases is represented by a distinct color. Dashed horizontal lines indicate the eventual extinction probability *q*(∞) for each disease.

To elucidate the influence of offspring distribution and generation time on the finite-time extinction probability, we calculated *T*_*reach*_, defined as the time required for *q*(*t*; *T*_*g*_) to reach *q*(∞) within a tolerance of 0.01 for various parameter combinations (**Fig. 5)**. For a fixed generation time *T*_*g*_, the basic reproduction number *R*_0_ exerts a significantly stronger influence on *T*_*reach*_ than does the dispersion parameter *k*. In particular, as *R*_0_ > 1 increases, *T*_*reach*_ decreases markedly, indicating that higher values of *R*_0_ shorten the time required for the extinction probability to converge toward its asymptotic value. Nonetheless, the effect of the dispersion parameter *k* becomes particularly pronounced when *R*_0_ is near one. In these scenarios, an increase in *k* leads to a notable extension in *T*_*reach*_, reflecting a sensitivity to the heterogeneity in secondary infections when transmission is barely sustaining an outbreak. Regarding the role of the generation time, *T*_*g*_ appears to primarily scale the distributed patterns of *T*_*reach*_. In other words, variations in *T*_*g*_ effectively “zoom in” or “zoom out” on the contour of the *T*_*reach*_ landscape without fundamentally altering its overall structure. This observation indicates that while *T*_*g*_ modulates the absolute time scales of convergence, the relative sensitivity of *T*_*reach*_ to *R*_0_ and *k* is preserved.

**Figure 5.**
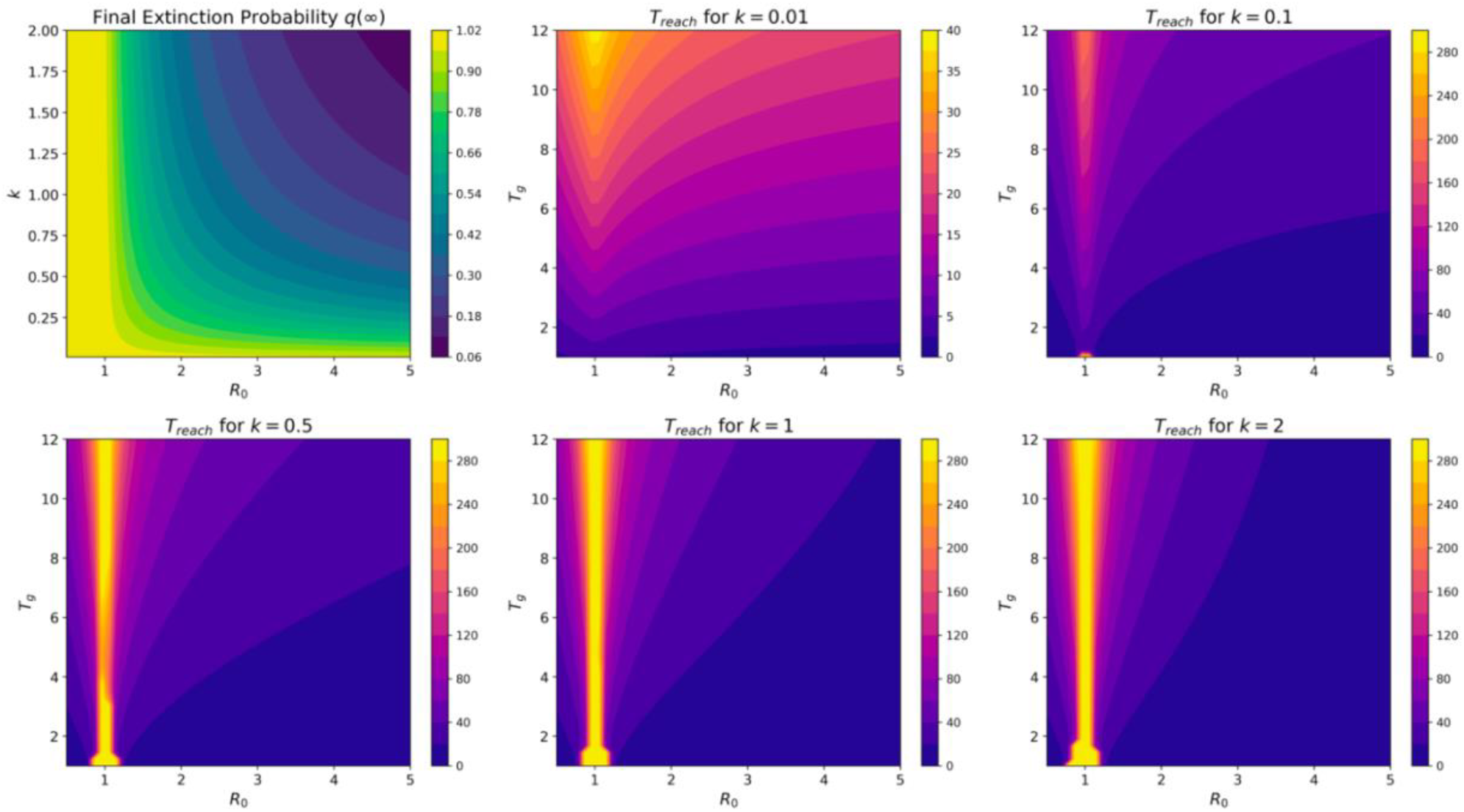
Composite figure with six subplots arranged in a 2 × 3 layout. The first panel presents a contour plot of the final extinction probability *q*(∞) as a function of the reproduction number *R*_0_ and the dispersion parameter *k*. The remaining five panels show contour plots of the time to reach extinction probability (*T*_*reach*_, within a tolerance of 0.01) as a function of *R*_0_ and mean generation time *T*_*g*_, for fixed values of *k* = 0.01, 0.1, 0.5, 2, and 10. Color bars in each subplot indicate the magnitude of the displayed quantity.

## 5 Discussion

Our study advances the understanding of epidemic extinction dynamics by incorporating the generation time distribution explicitly into the branching process framework. A key methodological contribution of this study is the explicit establishment of the finite-time extinction probability model by incorporating the generation time distribution. We comprehensively proved that the finite-time extinction probability, *q*(*t*; *T*_*g*_), decreases monotonically with respect to the mean time *T*_*g*_. This result not only provides a strong theoretical foundation for our model but also underscores the crucial role that timing plays in shaping the trajectory of epidemic fade-out.

The mathematical demonstration that *q*(*t*; *T*_*g*_) exhibits a monotone decline in relation to *T*_*g*_ is of significant importance. It confirms that, given an increase in the mean interval between transmission events, the extinction probability over any finite horizon is inherently lower. This counterintuitive insight highlights that while shorter generation times can accelerate the wave of infections and potentially hasten the early depletion of susceptible individuals, they promote a more rapid progression toward extinction, as the probability subsequently converges to its asymptotic value *q*(∞).

Our numerical simulations, supported by precise analytical proofs, further elucidate the interplay among the basic reproduction number *R*_0_, the dispersion parameter *k*, and the generation time *T*_*g*_. For instance, while an elevated *R*_0_ typically shortens the time required to approach *q*(∞), the dispersion parameter *k* plays a crucial role when *R*_0_ is close to one. These intricate dynamics illustrate that the temporal evolution of extinction probabilities is governed by a subtle balance between the average number of secondary infections and the inherent variability in the transmission process. In addition, the visual outcomes from our study reinforce these findings by demonstrating clear scaling effects driven by *T*_*g*_. Changes in the generation time predominantly act as a scaling mechanism for the extinction dynamics, effectively magnifying or contracting the overall timeline while leaving the underlying pattern dictated by *R*_0_ and *k* largely unaltered. This insight is particularly valuable as it implies that interventions aimed at modifying the generation time can systematically shift the temporal evolution of the epidemic.

Despite the robust nature of our model, it is crucial to recognize that our analysis relies on classical assumptions inherent in branching process frameworks, such as independent transmission events, a well-mixed population, and constant epidemiological parameters. Although these assumptions facilitate mathematical tractability and reliable numerical approximations, they represent an idealized version of pathogen spread. Future studies should expand the current model to more practical epidemiological settings by incorporating time-varying reproduction numbers and individual-level heterogeneities in generation time and offspring number. In addition, integrating realistic epidemic factors such as behavioral changes, network effects, and structured populations could provide in-depth insights into the mechanisms that drive epidemic persistence and fade-out in real-world scenarios. If certain diseases exhibit transmission behaviors that deviate from an exponential (memoryless) distribution, for instance, incubation and recovery times showing heavy-tailed or long-memory behavior, then integer-order models may struggle to capture these dynamics accurately. In such cases, fractional-order models offer a more flexible framework (Manivel and Kumawat 2025; Manivel et al. 2025; Shyamsunder et al. 2024; Shyamsunder and Purohit 2024; Shyamsunder 2025; Shyamsunder and Meena 2025; Soni 2025).

In conclusion, the finite-time extinction probability model we developed not only provides valuable quantitative insights into epidemic dynamics but also offers a crucial theoretical advancement by comprehensively proving the monotone decreasing property of *q*(*t*; *T*_*g*_) with respect to the mean generation time. These findings have crucial implications for public health policy, as they highlight the dual role of generation time: its reduction can accelerate early extinction, whereas its extension might delay the fade-out phase. Accordingly, public health interventions must weigh these considerations carefully to optimize strategies aimed at controlling infectious disease outbreaks.

## 6 Methods

The simulation is structured to estimate *q*(*t*; *T*_*g*_) by discretizing time and applying numerical integration to solve the renewal equation governing the process.

### 6.1 Model Parameterization

When visualizing the evolution of *q*(*t*; *T*_*g*_) with respect to time *t* and the mean generation time *T*_*g*_, the basic reproduction number *R*_0_ is set to 1.5. Two widely used offspring distributions— the Poisson and the geometric—are considered in the analysis that follows:

- **Poisson:** The PGF is defined by the following equation:

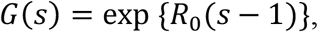

with no-offspring probability *p*_0_ = exp (−*R*_0_).
- **Geometric:** Given a success probability *p* = 1/(*R*_0_ + 1), the PGF is calculated as follows:

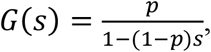

and *p*_0_ = *p*.

To capture individual variation, we use the negative binomial distribution parameterized by the mean *R*_0_ and dispersion *k*, which includes the Poisson distribution (*k* → ∞) and the geometric (*k* = 1) as special cases.

The generation time—defined as the delay between a primary infection and the occurrence of its secondary infections—is assumed to follow an exponential distribution, a common assumption in biological modeling due to its memoryless property. In particular, the generation time is modeled with the following density function and the CDF:

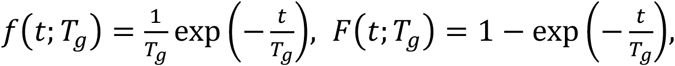

where *T*_*g*_ represents the mean generation time and sets the time scale of transmission events. For numerical integration, a discrete time grid over the interval [0, *T*_*max*_] with a fixed time step *dt* is constructed, providing the framework for approximating the integral in the renewal equation.

### 6.2 Numerical Solution Method

The evolution of the finite-time extinction probability *q*(*t*; *T*_*g*_) is governed by the renewal equation (Eq. (1)). To numerically solve this equation, we employ an iterative procedure based on the rectangular (Riemann sum) rule for integration. Starting with the initial condition *q*(0) = *p*_0_, the algorithm computes *q*(*t*; *T*_*g*_) at each discrete time step by approximating the integral as a summation over all preceding time points. This sequential updating across the discretized time domain yields the full temporal evolution of *q*(*t*; *T*_*g*_).

In parallel, to capture the long-term behavior of the system, the final asymptotic extinction probability *q*(∞) is determined by identifying the nontrivial fixed point of the governing equation

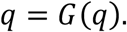

This fixed point is computed using a numerical root-finding method, particularly the *fsolve* function from SciPy to solve for *q*. The resulting probability serves as a benchmark, offering insight into the eventual fate of the branching process.

### 6.3 Calculation of the Time *T*_*reach*_ to Reach *q*(∞)

To quantify the timing of convergence of *q*(*t*; *T*_*g*_) to its asymptotic value *q*(∞), we define the time to reach the final extinction probability, *T*_*reach*_, as the earliest time point at which the absolute difference between the finite-time extinction probability and its asymptotic counterpart decreases below a prescribed tolerance (e.g., *ϵ* = 0.01). Formally, *T*_*reach*_ is determined by locating the smallest *t* that satisfies the following condition:

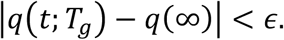

In practice, we evaluate |*q*(*t*; *T*_*g*_) − *q*(∞)| at each discrete time step and record the time *t* at which the condition is first satisfied. This approach provides a systematic means of capturing the convergence rate of the extinction probability and enables comparative analysis of how different parameter combinations—such as variations in *R*_0_, *k*, or *T*_*g*_ —influence the temporal dynamics of the epidemic. The metric *T*_*reach*_ serves as a critical indicator of the speed at which the process approaches its steady state, offering valuable insights for assessing intervention strategies and the inherent resilience of epidemic dynamics.

## Declarations

This work was supported by the National Natural Science Foundation of China (Grant Nos. 12301626), Guangdong Basic and Applied Basic Research Foundation (Grant Nos. 2022A1515110612), and Guangzhou Municipal Science and Technology Plan Basic and Applied Basic Research (Grant Nos. 2025A04J3668, 2025A03J3089). The funders had no role in the study design, data collection and analysis, decision to publish, or preparation of the manuscript.

## Code availability

All the codes used in the article can be found in Github: https://github.com/baoyinyuan/Extinction.git

## Author contribution

**Baoyin Yuan**: Conceptualization, Methodology, Data curation, Software, Visualization, Writing–original draft. **Feng Jiao**: Validation, Writing–review & editing, Funding acquisition.

## Appendix A

### Differentiability of *q*(*t*; *T*_*g*_) with Respect to *T*_*g*_

In this appendix, we prove that the finite-time extinction probability *q*(*t*; *T*_*g*_) depends continuously and differentiably on the parameter *T*_*g*_ under suitable smoothness conditions.

#### A.1 Notation and Assumptions

1. **Function Space**: Let *X* = *C*([0, *T*]) be the Banach space of continuous functions on [0, *T*] with the norm 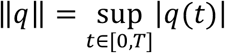
2. **Renewal Equation**: The extinction probability is given by

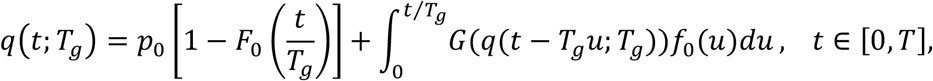

where 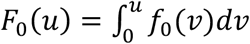 represents the cumulative distribution function associated with the waiting time density *f*_0_. The parameter *p*_0_ is a probability in the interval (0,1), and the functions *G*: ℝ→ ℝ and *f*_0_: [0, ∞) → ℝ are assumed to be at least *C*^1^(i.e., continuously differentiable) and bounded, with the additional property that *f*_0_(*u*) > 0 for all *u* ≥ 0. The parameter *T*_*g*_, which represents the mean generation time, is considered in a neighborhood around some nominal value *T*_*g*,0_.

#### A.2 Reformulation as a Fixed-Point Problem

Define the operator 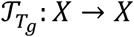 as

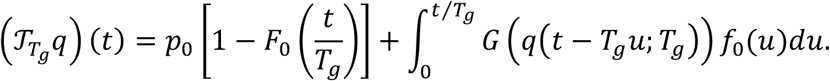

Then the renewal equation can be recast as a fixed-point equation:

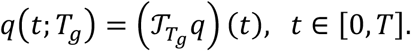

To analyze the dependence on *T*_*g*_, we define the mapping

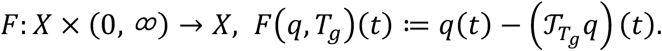

A solution *q*(.; *T*_*g*_) satisfies

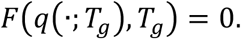

Our goal is to show that the mapping *T*_*g*_ ↦ *q*(.; *T*_*g*_) is continuously differentiable. This will be achieved by verifying that *F* is *C*^1^and that the Frechet derivative with respect to *q* at a solution point is invertible. Then the Banach Implicit Function Theorem applies.

#### A.3 Differentiability of *F*

##### A.3.1 Differentiability with Respect to *q*

Let *q* ∈ *X* and *v* ∈ *X* be an increment. We have

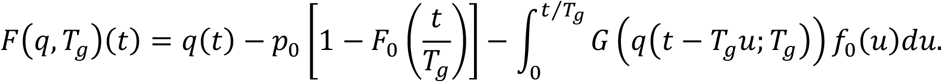

Note that the first term, *q*(*t*), is linear in *q*. The term *p*_0_[1 − *F*_0_(*t*/*T*_*g*_)] does not depend on *q*. In the integral term, since *G* is *C*^1^, applying the chain rule we have

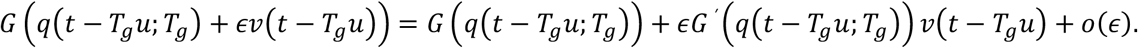

Thus, the Fréchet derivative of *F* in the direction *v* is given by

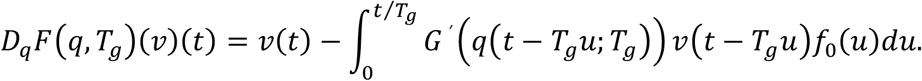

It is convenient to express the Fréchet derivative with respect to *q* as

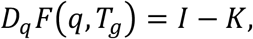

where *I* denotes the identity operator on the space *X* = *C*([0, *T*]) (that is, (*Iv*)(*t*) = *v*(*t*) for all *v* ∈ *X* and *t* ∈ [0, *T*]), and the linear operator *K* is defined by

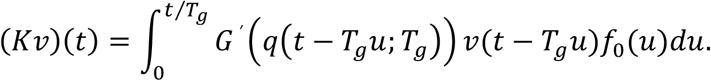

##### A.3.2 Differentiability with Respect to *T*_***g***_

The parameter *T*_*g*_ appears in several places: in the scaling *t*/*T*_*g*_ inside the function *F*_0_, as the upper limit of the integral, in the time shift *t* − *T*_*g*_*u*, and also in the implicit dependence of *q*(*t* − *T*_*g*_*u*; *T*_*g*_). Differentiability with respect to *T*_*g*_ is to be studied by applying the chain rule and Leibniz’s rule. Specifically, for the term *p*_0_[1 − *F*_0_(*t*/*T*_*g*_)], we have

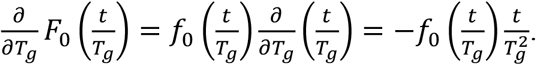

Similarly, for the integral term, we apply Leibniz’s rule. Although the derivative

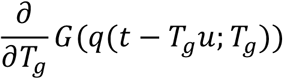

formally yields

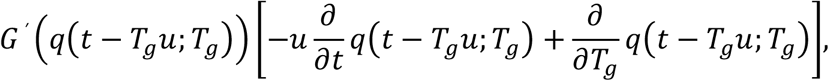

the existence of this derivative ultimately follows from applying the implicit function theorem, which guarantees that the solution *q*(.; *T*_*g*_)is differentiable with respect to *T*_*g*_. Since our focus is solely on establishing the differentiability of *F* with respect to *T*_*g*_ rather than determining its explicit form, we assume under our hypothesis that all necessary derivatives exist, thereby justifying the formal application of the chain rule.

#### A.4 Invertibility of *D*_*q*_*F*(*q, T*_*g*_)

The next crucial step is to show that the Fréchet derivative with respect to *q*,

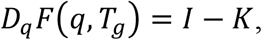

is an isomorphism on *X*.

##### A.4.1 Estimating the Operator *K*

For any *v* ∈ *X*, we have

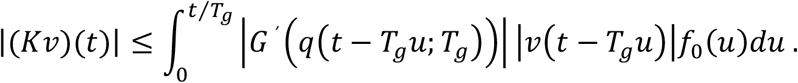

Let

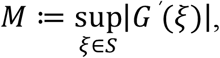

where *S* is a compact set containing the range of *q*(*t*; *T*_*g*_). Then, since |*v*(*t* − *T*_*g*_*u*)| ≤ ‖*v*‖_∞_, it follows that

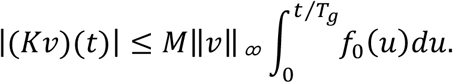

Taking the supremum over *t* ∈ [0, *T*] gives

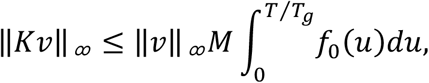

so that the norm of *K* satisfies

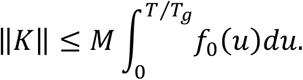

##### A.4.2 Sufficient Condition for Invertibility

A sufficient condition for *I* − *K* to be invertible is that

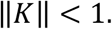

This condition can be guaranteed by one or more of the following: (i) Small observation time (*T*): choosing *T* sufficiently small makes the integration interval [0, *T*/*T*_*g*_] small, thereby reducing the value of the integral; (ii) Bound on *G*^′^ : if *G* is sufficiently smooth such that its derivative *G*^′^ is bounded by a small constant *M*, the norm reduction follows; (iii) Small integral of *f*_0_: if the waiting-time density *f*_0_(*u*) is low on the interval [0, *T*/*T*_*g*_], for example, if the event that triggers reproduction or change occurs slowly—then the overall integral remains small.

Under these assumptions, we have ‖*K*‖ < 1, and consequently, the 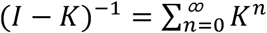 converges. Hence, *I* − *K* is invertible.

#### A.5 Application of the Implicit Function Theorem

Since we have shown that the mapping *F*: *X* × (0, ∞) → *X*, 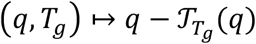 is continuously differentiable (*C*^1^) with respect to both *q* and *T*_*g*_, and the Fréchet derivative with respect to *q* at a solution(*q*_0_, *T*_*g*,0_) (i.e., *F*(*q*_0_, *T*_*g*,0_) = 0), *D*_*q*_*F*(*q*_0_, *T*_*g*,0_) = *I* − *K* is invertible by the estimate in the above section A.4.

By the Banach Implicit Function Theorem, there exists a unique local mapping *T*_*g*_ ↦ *q*(.; *T*_*g*_) in a neighborhood of *T*_*g*,0_ such that *F*(*q*(.; *T*_*g*_), *T*_*g*_) = 0. Moreover, the mapping *T*_*g*_ ↦ *q*(.; *T*_*g*_) is continuously differentiable. In particular, the partial derivative

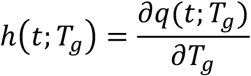

exists and is continuous on the considered interval.

**End of Appendix A**.

## Appendix B

### Derivation of Finite-Time Extinction Probability to Include Mean Generation Time

In this Appendix, we provide a step-by-step derivation of the finite-time extinction probability formula and show that its derivative with respect to the mean generation time,

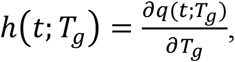

satisfies a Volterra-type integral equation.

#### B.1 Construction of the Finite Extinction Probability Formula

We assume that the discrete offspring distribution of an individual is characterized by the generating function

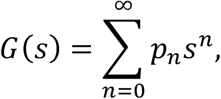

With

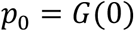

being the probability that the individual produces no offspring.

Let the generation (or reproduction) time have a density 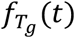 and corresponding cumulative distribution function

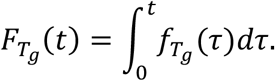

Under the assumption of a scaling family, these functions can be written as

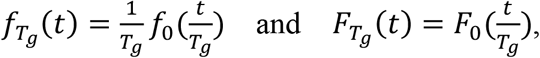

where *f*_0_ and *F*_0_ are the standard density and cumulative distribution functions, respectively.

The finite-time extinction probability *q*(*t*; *T*_*g*_) is defined as the probability that the initial individual and all of its descendants are extinct by time *t*. Conceptually, two mutually exclusive scenarios are considered:

1. **No reproduction occurs in** [**0, *t***]: The probability that the reproduction time exceeds *t* is

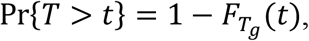

and extinction occurs if the individual produces no offspring. Hence, the contribution is

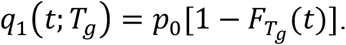
2. **At least one reproduction occurs in** [**0, *t***]: Let the first reproduction occur at time *τ* ∈ [0, *t*] with density 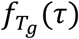.Upon reproduction, suppose the individual produces *n* ≥ 0 offspring (with probability *p*_*n*_). Each descendant, independently, goes extinct by time *t* with probability *q*(*t* − *τ*; *T*_*g*_). Averaging over *n* (using the generating function), the conditional extinction probability is

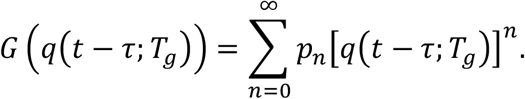

Integrating over all reproduction times *τ* gives

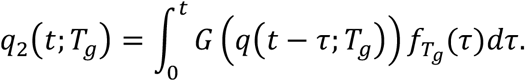

Thus, by the law of total probability, the overall finite-time extinction probability is

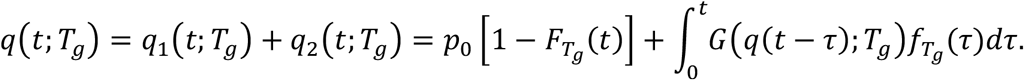

Using the scaling expression for 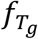 and 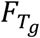, we rewrite the above as

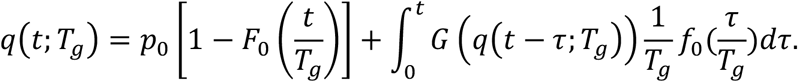

Changing the variable via 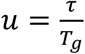 (so that *τ* = *T*_*g*_ *u* and *dτ* = *T* _*g*_ *du*, the update equation becomes

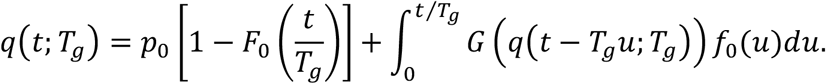

#### B.2 Derivation of the Volterra-type Integral Equation for the Derivative

We now differentiate *q*(*t*; *T*_*g*_) with respect to *T*_*g*_ and show that the derivative

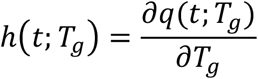

satisfies a Volterra-type integral equation.

**Step B.2.1: Differentiation of the First Term**. The first term is

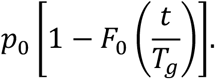

Differentiating with respect to *T*_*g*_ via the chain rule,

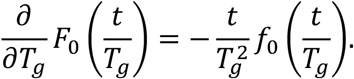

Thus, the derivative of the first term is

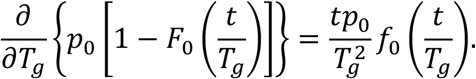

**Step B.2.2: Differentiation of the Integral Term**. The integral term is given by

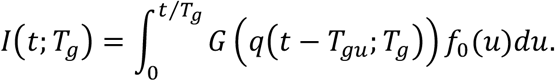

Differentiation involves two contributions: one from the upper limit and one from differentiating the integrand.

- *Upper Limit Contribution:* The upper limit is *t*/*T*_*g*_. Its derivative is

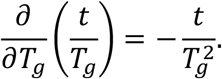

At *u* = *t*/*T*_*g*_, note that

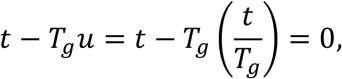

and since *q*(0; *T*_*g*_) = *p*_0_, we have

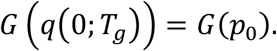

Consequently, the contribution is

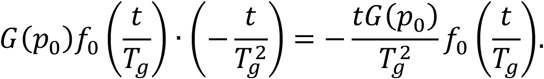

- *Differentiation Under the Integral Sign:*

Differentiate the integrand with respect to *T*_*g*_. By the chain rule,

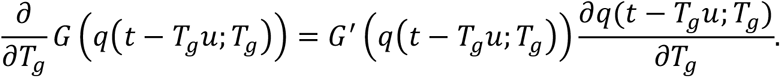

Define

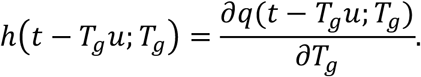

The corresponding contribution is

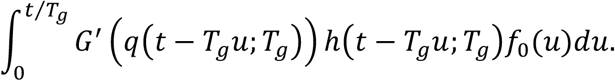

**Step B.2.3: Combining Contributions**. Summing the derivatives of the first and integral parts, we obtain

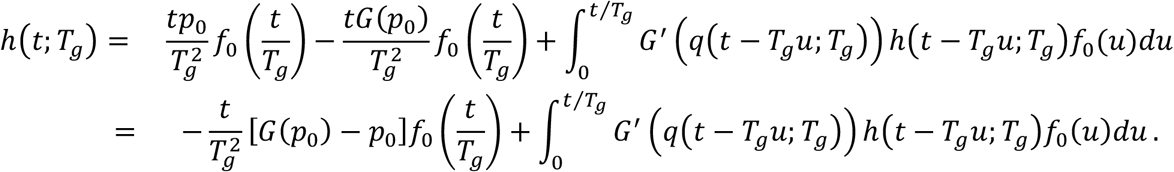

This is the desired Volterra-type integral equation for *h*(*t*; *T*_*g*_).

**End of Appendix B**.

## Appendix C

### On the Strict Negativity of the Sensitivity of Finite-Time Epidemic Extinction to Mean Generation Time

In this appendix we provide a complete and detailed proof that the partial derivative

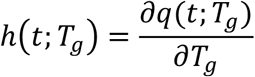

of the finite-time extinction probability *q*(*t*; *T*_*g*_) with respect to the mean generation time *T*_*g*_ is strictly negative for all *t* > 0. The proof proceeds as follows:

i. We introduce an auxiliary function to recast the integral equation for *h*(*t*; *T*_*g*_) into one that is amenable to fixed-point iteration.
ii. We then construct an iteration scheme and prove that the associated operator is a contraction using the Banach fixed-point theorem.
iii. We establish the convergence, uniqueness of the fixed point, and finally extend this local result to the entire finite interval.

#### C.1 Construction of the Auxiliary Function

Recall that the derivative *h*(*t*; *T*_*g*_) satisfies the Volterra-type integral equation

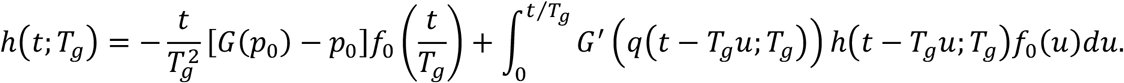

For convenience, we introduce the auxiliary function

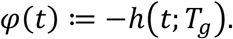

Thus, the above equation becomes

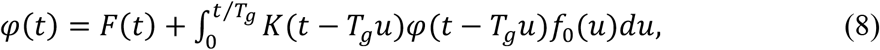

Where

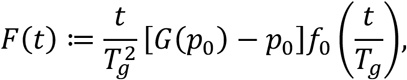

And

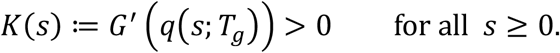

It is important to note that, since the offspring distribution is assumed to be nontrivial (i.e., *p*_0_ < 1 and at least one *p*_*k*_ > 0 for some *k* ≥ 1), we have 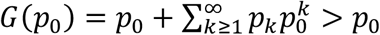.Thus, *G*(*p*_0_) − *p*_0_ > 0, which immediately implies that *F*(*t*) > 0 for all *t* > 0 (given that *f*_0_(*t*/*T*_*g*_) > 0).

Showing that *φ*(*t*) > 0 for all *t* > 0 is equivalent to proving *h*(*t*; *T*_*g*_) = −*φ*(*t*) < 0, which is the desired monotonicity result.

#### C.2 Iterative Fixed-Point Construction and Contraction Mapping

Define an operator *T* on a suitable function space, for example, the space of continuous functions *C*([0, *T*]) on a finite interval [0, *T*] (with the supremum norm ‖.‖_∞_ and *T* > 0 can be any finite time), by

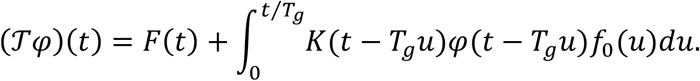

We wish to show that *T* is a contraction mapping on a closed subset of *C*([0, *T*]).

**Step C.2.1: Iteration Scheme**. Start with the initial function

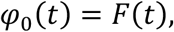

and define the iterative sequence

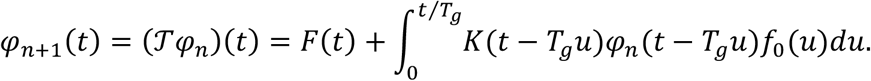

Since *F*(*t*) > 0 for all *t* > 0 and the integrand is strictly positive (due to *K*(.) and *f*_0_(*u*) > 0), we have

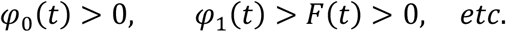

Thus, by induction, *φ*_*n*_(*t*) > 0 for all *n* and for all *t* ∈ [0, *T*].

**Step C.2.2: Proving the Contraction Property**. Let *φ, ψ* ∈ *C*([0, *T*]). Then

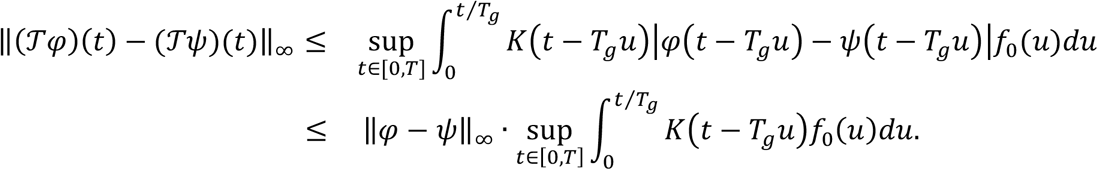

Taking the supremum over *t* ∈ [0, *T*] yields

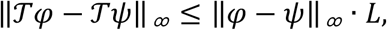

with

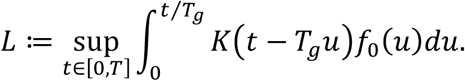

For *T* to be a contraction, we require that *L* < 1. Note that the contraction constant *L* depends on three key components. First, it is influenced by the length of the observation interval [0, *T*]. Since the integration is carried out over *u* ∈ [0, *t*/*T*_*g*_] with *t* ≤ *T*, the maximum integration length is *T*/*T*_*g*_. By choosing *T* sufficiently small, or by considering the analysis over a local interval, it is possible to ensure that the integral 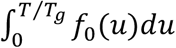 is small. Second, the bound on the kernel *K*(*s*) = *G*^′^(*q*(*s*; *T*_*g*_)) plays an important role. If we assume there exists a constant *M* > 0 such that 0 < *K*(*s*) ≤ *M* for all *s* ∈ [0, *T*], then it follows that 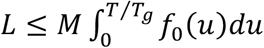. Thus, one obtains a sufficient condition for the contraction given by 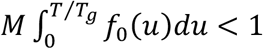. Third, the behavior of the waiting-time density *f*_0_(*u*) is crucial. If the waiting-time density *f*_0_(*u*) is small on the integration interval—for instance, if events occur with relatively low instantaneous probability—then the overall integral will remain limited.

Under the above conditions, i.e., either by choosing small *T* relative to *T*_*g*_ or by ensuring that 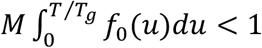 via additional structural properties of the model, the operator *T* becomes a contraction mapping on *C*([0, *T*]).

**Step C.2.3: Application of the Banach Fixed-Point Theorem**. Since *T* is a contraction on the complete metric space *C*([0, *T*]), by the Banach Fixed-Point Theorem, there exists a unique fixed point *φ*^∗^(*t*) ∈ *C*([0, *T*]) such that

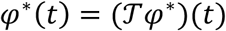

for all *t* ∈ [0, *T*]. Moreover, the iterative sequence {*φ*_*n*_(*t*)} converges uniformly (with respect to *t*) to *φ*^∗^(*t*).

#### C.3 Uniqueness, Convergence and Extension (Continuation)

**Uniqueness and Convergence:** The uniqueness of the fixed point implies that the solution *φ*(*t*)of Equation (8) is unique on the interval [0, *T*]. By the estimate in Step B.2.2, the error after *n* iterations is bounded by

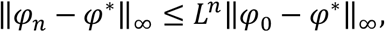

which shows uniform convergence as *n* → ∞.

**Extension (Continuation) to the Whole Finite Interval:** The above contraction mapping argument applies on a small interval [0, *T*_0_] where *T*_0_ is chosen so that the contraction constant *L* < 1. To extend the result to any *T* > 0, one partitions the interval [0, *T*] into a finite number of subintervals [0, *T*_0_], [*T*_0_, 2*T*_0_], ⋯ and applies the local result in each subinterval. The continuity and uniqueness of the fixed points on the overlapping endpoints ensure that the local solutions can be concatenated to yield a unique solution *φ*(*t*) > 0 for all *t* ∈ [0, *T*].

#### C.4 Conclusion of the Monotonicity Proof

Since the unique fixed-point solution *φ*(*t*) obtained via the iteration satisfies

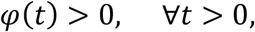

and recalling the definition

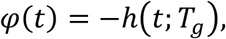

we conclude that

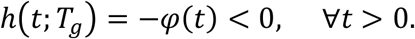

This completes the proof that the partial derivative of \*q*(*t*; *T*_*g*_) with respect to *T*_*g*_ is strictly negative, i.e., the finite-time extinction probability is strictly decreasing with respect to the mean generation time.

**End of Appendix C**.

